# Microbiota-dependent elevation of Alcohol Dehydrogenase in *Drosophila* is associated with changes in alcohol-induced hyperactivity and alcohol preference

**DOI:** 10.1101/444471

**Authors:** Malachi A. Blundon, Annie Park, Scott A. Keith, Stacie L. Oliver, Rory A. Eutsey, Anna M. Pyzel, Tiffany W. Lau, Jennifer H. Huang, Hannah M. Kolev, N. Luisa Hiller, Nigel S. Atkinson, Jonathan S. Minden, Brooke M. McCartney

**Affiliations:** Department of Biological Sciences, Carnegie Mellon University, Pittsburgh, PA 15213, USA; Department of Neuroscience and Waggoner Center for Alcohol and Addiction Research, University of Texas at Austin, Austin, TX 78712, USA

**Keywords:** microbiota, 2D-DIGE, alcohol dehydrogenase, alcohol, *Drosophila*, alcohol induced hyperactivity, alcohol preference

## Abstract

The gut microbiota impacts diverse aspects of host biology including metabolism, immunity, and behavior, but the scope of those effects and their underlying molecular mechanisms are poorly understood. To address these gaps, we used Two-dimensional Difference Gel Electrophoresis (2D-DIGE) to identify proteomic differences in male and female *Drosophila* heads raised with a conventional microbiota and those raised in a sterile environment (axenic). We discovered 22 microbiota-dependent protein differences, and identified a specific elevation in Alcohol Dehydrogenase (ADH) in axenic male flies. Because ADH is a key enzyme in alcohol metabolism, we asked whether physiological and behavioral responses to alcohol were altered in axenic males. Here we show that alcohol induced hyperactivity, the first response to alcohol exposure, is significantly increased in axenic males, requires ADH activity, and is modified by genetic background. While ADH activity is required, we did not detect significant microbe-dependent differences in systemic ADH activity or ethanol level. Like other animals, *Drosophila* exhibit a preference for ethanol consumption, and here we show significant microbiota-dependent differences in ethanol preference specifically in males. This work demonstrates that male *Drosophila’s* association with their microbiota affects their physiological and behavioral responses to ethanol.

## Introduction

The human microbiota, the community of microorganisms including bacteria and fungi that resides in and on our bodies, contribute to metabolism, immunity, and defense against pathogens [1], [2]. Surprisingly, recent evidence suggests that the bacterial microbiota of the gut can also influence learning, memory, anxiety, depression, and autism-associated behaviors in some animals [3]–[6]. The number of connections being made between symbiotic bacteria and host physiologies and behaviors is rapidly increasing, making it likely that more associations await discovery. Furthermore, we understand relatively little about the molecular mechanisms that mediate any of these host-microbe interactions.

*Drosophila* is emerging as an excellent model to dissect the role of the microbiota in animal physiology and behavior. Bacteria in the order *Lactobacillales* are found in both the *Drosophila* and human microbiota [7]–[9], and links between the microbiota and host physiology and behavior are also present in *Drosophila*. Fly behaviors such as egg laying, feeding, male competition, and kin recognition all respond to changes in the microbiota [10]–[18]. The fly microbiota can modulate insulin, insulin-like growth factor, and Target Of Rapamycin (TOR) signaling thereby affecting systemic homeostasis in the fly [19], [20]. Furthermore, host-pathogen studies in *Drosophila* have proven invaluable to unraveling the mechanisms of human innate immunity [21], [22]. Thus, *Drosophila* provides an excellent model for host-microbe interactions.

Proteome analysis provides valuable information about protein abundance and post-translational modifications (PTMs) that can be missed at the transcriptome level [14]–[17]. Importantly, it has been shown that there is little correlation between mRNA expression and protein abundance [23]–[27]. While several studies have focused on microbe-dependent transcriptome changes in the *Drosophila* gut or in the whole fly [28]–[34], no proteomic analysis has been done. Two-Dimensional Difference Gel Electrophoresis (2D-DIGE) is a powerful technique to reveal proteomic changes between two or three protein samples simultaneously run on the same gel [35]–[38]. Protein differences detected by 2D-DIGE are then identified using liquid chromatography coupled to tandem mass spectrometry (LC-MS/MS).

Here we used 2D-DIGE to identify *Drosophila* proteins that are responsive to the microbiota. We focused on the *Drosophila* head proteome to search for proteins with potential roles in neural function and behavior, as this aspect of host-microbe interactions is not well understood. By comparing the head proteomes of male or female flies raised with a conventional microbiota (CV) to those raised in a sterile environment (axenic, AX), we identified 22 proteins with altered abundance. Interestingly, several of these differences were sex specific. One of the male-specific difference-proteins is Alcohol Dehydrogenase (ADH), a key enzyme in ethanol metabolism in all animals, which was increased in AX males and reversed by reintroducing the conventional microbiota. ADH elevation suggested that AX males may have altered physiological and behavioral responses to alcohol. Indeed, we found that AX males exhibited significantly enhanced alcohol-induced hyperactivity (AIH), a response that is ADH dependent, male specific, and sensitive to host genetic background and dietary conditions. Using different measures of ethanol preference, we found that when offered a choice, AX males preferred to consume food containing alcohol significantly more than their CV siblings. Taken together, our work demonstrates a novel connection between the microbiota and host physiological and behavioral responses to alcohol in *Drosophila* that may have implications for our understanding of the microbiota’s role in alcohol use disorders (AUD).

## Results

### The *Drosophila* head proteome is responsive to microbial condition

To identify proteome changes in the heads of *Drosophila* with or without gut microbiota, we compared head lysates (Fig. 1A) of CV and AX siblings (SFig. 1) from a wild type strain called “Top Banana” (see methods). Lysates from CV and AX fly heads were independently labeled with either Cy3 or Cy5 2D-DIGE dyes. The labeled protein lysates were then combined and run on the same 2D-DIGE gel (Fig. 1A). We detected and quantified the protein spots using an open-source astronomy software package called SourceExtractor, as previously described [35], [36], [39]. Because we saw variability in host protein expression similar to what has been observed in host microbiota-dependent mRNA expression studies [40], [41], we used two criteria to determine whether a protein was different in CV vs AX heads. First, we used a 20% difference in protein abundance (1.2-fold) as a cut off for a significant protein expression difference; this is approximately three standard deviations above the technical noise in a standard 2D-DIGE experiment [36]. Second, significant proteins must be different in at least two of the three biological replicates. Using these criteria, we identified 22 and 16 reproducible difference-proteins in male and females, respectively (Fig. 1B,C, SFig. 2A&B). All of these difference-proteins exhibited protein abundance changes. While some of the differences were shared between males and females, most appeared to be sex-specific: four were male-specific (Fig. 1B&C, blue shaded region) and six were female specific (Fig. 1B&C, grey shaded region). Among the shared proteins, three changed abundance in the same direction in AX male and female flies (Fig. 1B&C, orange shaded region) and 3 changed abundance in the opposite direction (Fig. 1B&C, red shaded region). Additionally, six protein differences detected in the male head gels did not resolve in the female head gels (Fig. 1B, “did not compare” group). Together, our 2D-DIGE analyses revealed microbiota-dependent sexually dimorphic changes in the *Drosophila* head proteome.

**Figure 1.**
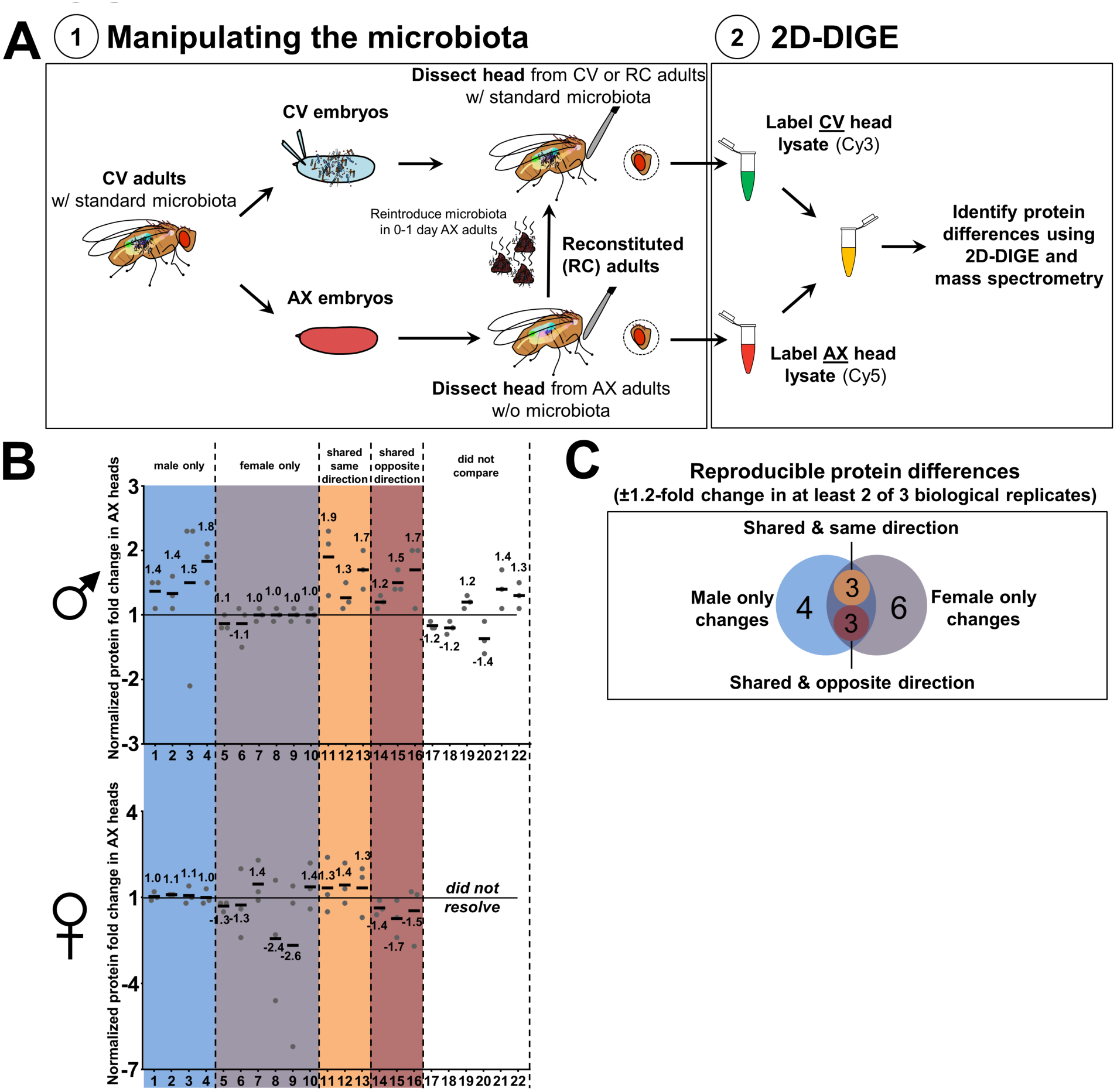
Identification of microbe-dependent differences in Drosophila male and female head proteomes. **(A)** Manipulating the microbiota and 2D-DIGE screen experimental design. (1) CV, AX, and RC siblings were derived from cohorts of embryos with the same parents. CV embryos were not manipulated, while AX embryos were dechorinated and reared under sterile conditions (see methods). 0-1 day old CV and AX adult flies were transferred to vials and aged for 5-6 days. For microbial reconstitution (RC), 0-1 day old AX flies were transferred to vials conditioned with feces from CV males and aged for 5-6 days. (2) Protein lysates were prepared from dry dissected heads, covalently labeled with either propyl-Cy3 or methyl-Cy5, combined, and co-electrophoresed on a 2DE gel. Difference proteins were identified by LC-MS/MS. **(B)** Measured fold-change for each protein difference was calculated using the Cy3 and Cy5 raw fluorescence intensities. Each dot represents a biological replicate containing 40 fly heads. Solid black bars indicate the mean. Six difference proteins identified in males (17-22) could not be resolved in the gels with female samples. **(C)** The Venn diagram shows a summary of the 16 reproducible protein differences for male and female data sets across three biological replicates. The color code matches that of panel B.

### Alcohol Dehydrogenase protein level is elevated in the heads of AX male flies

LC-MS/MS identified spot #4 (SFig. 2A&B) as the metabolic enzyme Alcohol Dehydrogenase (ADH). We confirmed the protein identity by immunoblotting for *Drosophila* ADH after 2DE separation of CV head lysate (Fig. 2A). 2D-DIGE analysis showed that the loss of microbiota leads to elevated ADH protein in the heads of AX males, but not AX females (Fig. 1B and 2B). On average, ADH protein was elevated 1.8-fold in AX males. To confirm that the microbiota influences ADH protein levels, microbes were reintroduced to 0-1 day old AX adult males by exposure to CV fecal deposits (referred to as Reconstituted flies, RC; Fig. 1A). ADH protein levels were restored to CV levels in RC males (Fig. 2B). One potential mechanism for the elevation of ADH protein is an increase in gene expression, but we did not find a consistent elevation of *Adh* transcripts in either AX male or female heads (Fig. 2C). Previous work had shown that ADH is not significantly expressed in fly brain [42], and consistent with this we find detectable ADH protein in the head capsule only (the head tissue without the brain; Fig. 2D). Because high levels of ADH are expressed in the abdominal fat body [42], the most likely site of ADH expression in the head is in the fat body that lies immediately anterior to the brain. Together, these data suggest a model in which the microbiota affect the level of ADH protein in the male head, likely in the fat body, through a mechanism that regulates ADH protein stability or translation efficiency.

**Figure 2.**
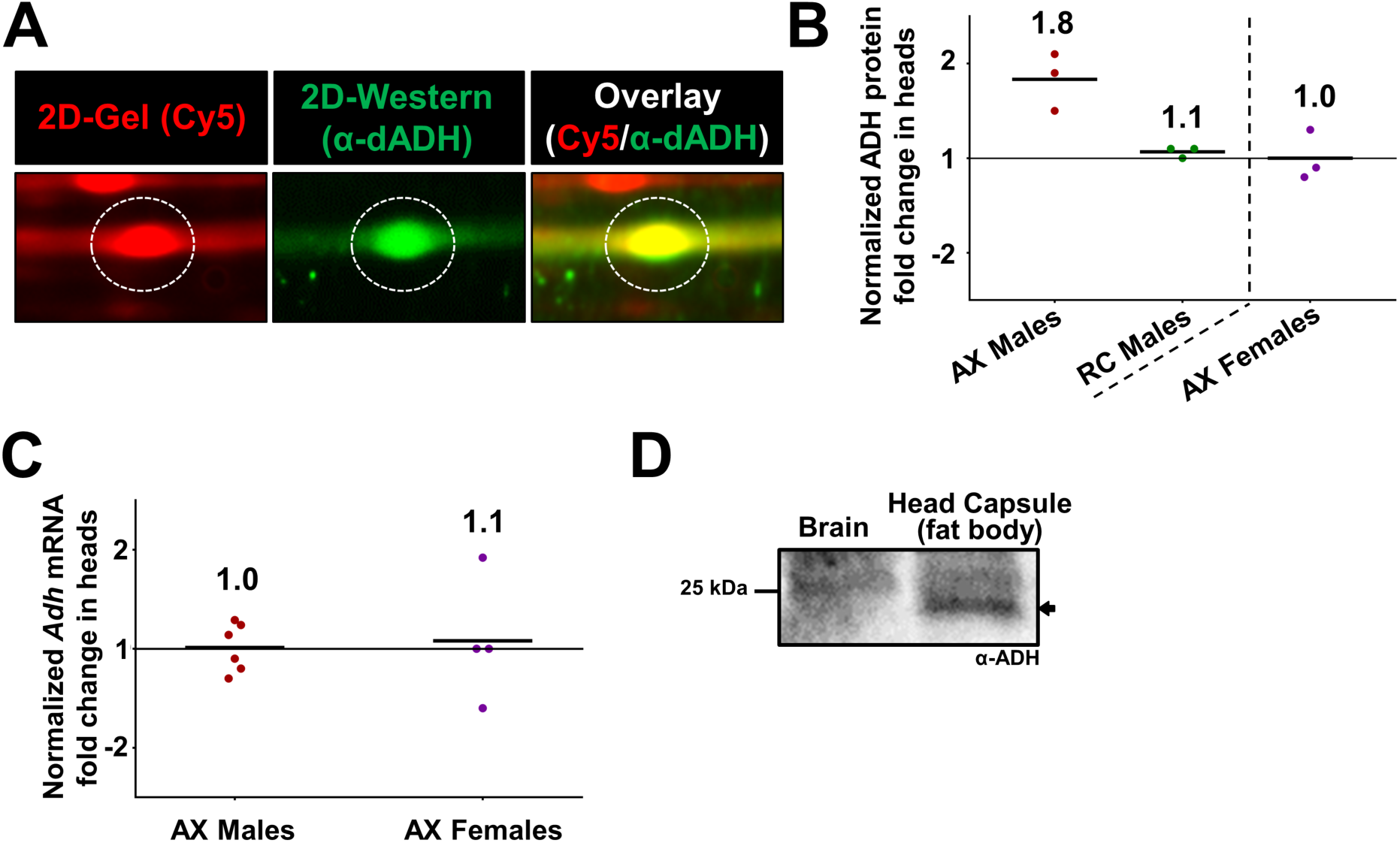
Alcohol Dehydrogenase (ADH) protein, but not mRNA, is elevated in AX male heads and decreased with microbial reconstitution. **(A)** 2D-Western blot of the ADH protein in the CV male head proteome. The overlay of the Cy5 labeled protein (red) with the signal from the anti-ADH immunoblot (green) confirmed the protein identification made by LC-MS/MS. **(B)** Reconstitution (RC) of the microbiota in AX adult males restored ADH to CV levels. AX male and female data shown were re-plotted from Fig. 1B. **(C)** RT-qPCR revealed no change in *Adh* transcription in AX compared to CV heads when normalized to *rpl32*. **(D)** Anti-ADH immunoblot of isolated CV male brains and head capsules (containing fat body) shows detectable ADH only in the head capsule (arrow).

The increase in ADH protein could result in increased ADH activity and increased ethanol metabolism. To test this, we quantified ADH enzymatic activity and ethanol levels in CV, AX, and RC males after exposing them to ethanol vapor. Because of assay limitations, it was not technically feasible to do these measurements with only heads. We thus characterized whole fly ADH activity and ethanol metabolism. These assays did not indicate any significant microbiota-dependent differences (SFig. 3A,B). While this does not rule out the possibility of tissue or cell type specific increases in ADH activity and ethanol metabolism, it does indicate that there is no significant systemic change.

### AX male flies are more responsive to alcohol

ADH catalyzes the oxidation of ethanol to acetaldehyde, which is the first step of ethanol metabolism. Thus, ADH influences several physiological responses to ethanol, including locomotor-hyperactivity [43]–[45]. Given the elevated ADH protein in AX male fly heads, we predicted that they may exhibit altered physiological responses to alcohol. To test this, we assessed alcohol induced hyperactivity (AIH) and sedation, two phases of alcohol induced responses common to all animals and well described in *Drosophila* [46], [47].

To assess AIH, we monitored locomotor activity of CV, AX, and RC males using the *Drosophila* activity monitor 2 (DAM2). This automated system uses infrared beams to quantify fly motility in the absence or presence of ethanol (Fig. 3A). Because ADH activity is required for AIH (SFig. 4A; [45]), we reasoned that AX males could have increased AIH due to elevated ADH protein. After monitoring baseline locomotion for 60 minutes, we exposed the flies to a low concentration of ethanol vapor (10:1 air to ethanol vapor) and continued monitoring for 120 minutes. Shortly after being exposed to ethanol vapor, CV males entered a period of hyperactivity peaking at an average of 3.4 beam passes/10 min (Fig. 3B). AX males entered the hyperactivity phase during the same time frame with a peak of 9.5 laser passes/10 min. This was restored to CV levels with microbial reconstitution in RC males (peak of 3.4 laser passes/10 min; Fig. 3B). Consistent with the lack of ADH elevation in AX female heads, there was no difference in AIH between CV and AX females (SFig. 4B).

**Figure 3.**
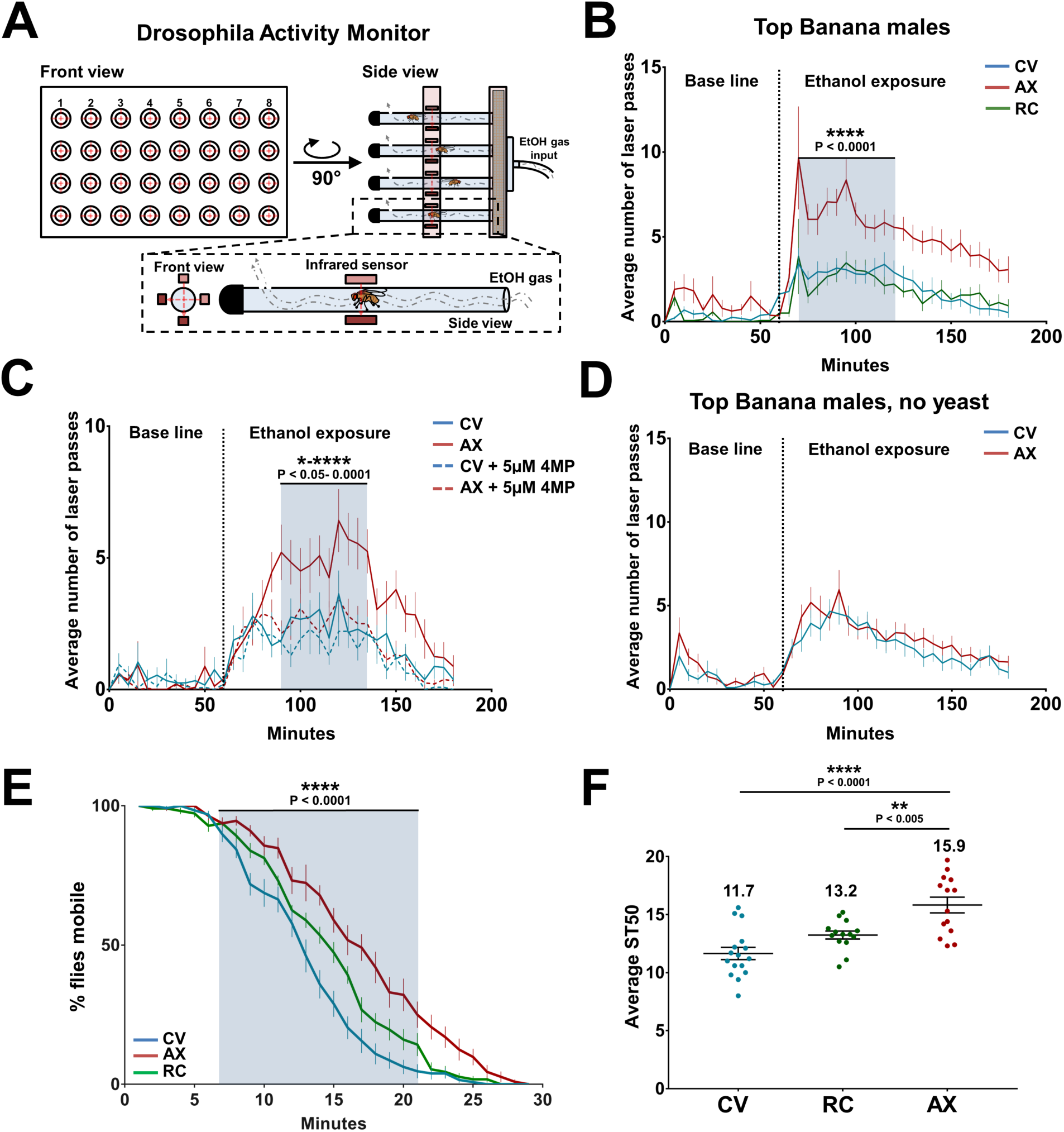
Microbe-dependent differences in hyperactivity and sedation responses to alcohol vapor exposure. **(A)** A schematic of the *Drosophila* Activity Monitor used to test alcohol induced hyperactivity (AIH). **(B-D)** In all experiments examining AIH, the flow ratio was H_2_O:EtOH (10:1) and the dotted lines indicate start of ethanol exposure. The error bars indicate standard error of the mean and statistical significance (blue shaded region) was assigned by a two-way ANOVA using **(B)** Tukey’s multiple comparisons post hoc test or **(C)** Dunnett’s multiple comparisons post hoc test. **(B)** AX males exhibited significantly elevated AIH compared to their CV and RC siblings. n = 32 flies in 4 trials/condition. **(C)** AIH in CV and AX males with and without 5 μM 4-Methylpyrazole demonstrated that ADH activity is necessary for the elevated AIH in AX males. n = 24 flies in 3 trials/condition. **(D)** AIH in CV and AX males raised in the absence of autoclaved inactive yeast granules shows that nutrient deprivation suppresses the elevated AIH in AX males. n = 32 flies in 4 trials/condition. **(E-F)** AX males exhibit delayed alcohol induced sedation compared to their CV and RC siblings. **(E)** Alcohol induced sedation curves for CV (n=128 in 16 vials), AX (n=112 in 14 vials), and RC (n=112 flies in 14 vials) males averaged from 4 trials. Bars indicate standard error of the mean. Statistical significance (blue shaded region, CV vs. AX comparison) was assigned by a two-way ANOVA (Dunnett’s multiple comparisons post hoc test). **(F)** Average ST50 derived from the alcohol induced sedation curves in panel E. Each dot represents the ST50 for a single vial. Bars indicate standard error of the mean and statistical significance was assigned by a one-way ANOVA (Tukey’s multiple comparisons post hoc test).

To understand the connection between increased ADH protein and AIH elevation, we asked whether AX males require ADH activity for their elevated AIH by treating CV and AX males with an ADH inhibitor, 4-Methylpyrazole [48]. If the elevated AIH in AX males requires ADH enzymatic activity, inhibiting ADH should reduce hyperactivity to the lower levels observed in CV. Indeed, AX males aged in the presence of the inhibitor for five days exhibited reduced AIH compared to control untreated AX males (Fig. 3C). Interestingly, the same inhibitor treatment of CV males did not significantly decrease their AIH, suggesting that the inhibitor treatment did not completely abolish ADH activity, or that there is a component of AIH that is ADH-independent (Fig. 3C).

Because diet and host genetic background are important for alcohol induced physiological changes, and influence the microbiota and its downstream effects [43], [49]–[53], we asked whether these factors influence microbiota-dependent AIH. First, we decreased protein availability to the adults by removing the autoclaved yeast supplement and found that this completely abolished the elevated AIH in AX males (Fig. 3D). To test if genetic background influences microbiota-dependent AIH, we examined AIH in two additional wild type lab strains, Canton S and Oregon R. Interestingly, there was no difference in AIH between Canton S CV and AX males (SFig. 4C). We did observe a significant increase in AIH in Oregon R AX compared to CV males (SFig. 4D), but the magnitude was less than what we found in Top Banana males (Fig. 3B). These data support the idea that diet and host genetic background interact with the microbiota to influence AIH in male flies.

Next, we asked whether the microbiota influences alcohol induced sedation, a physiological response that is largely independent of ADH activity [54]–[56]. To test this, we exposed groups of CV, AX, and RC males to ethanol vapor in fly vials and assessed the time to immobilization for the population [57]. The rate of sedation, assessed by comparing the time at which 50% of the population was immobilized (ST50), differed significantly between CV and AX males (Fig. 3E,F). AX males had a significantly higher ST50 (ST50=15.9 min) compared to CV males (ST50=11.7 min), and this was restored to CV levels in RC males (ST50=13.2 min). Overall, these data demonstrate that males have microbiota-dependent changes to AIH and sedation, and that the elevation in AIH requires ADH activity.

### AX males flies exhibit altered alcohol preference

Human studies have shown that differences in physiological responses to alcohol can influence alcohol consumption behavior and are a predictor of future alcohol addiction [58]. To determine whether the microbiota affects alcohol consumption and preference, we asked whether AX males have an altered alcohol preference. We first assessed alcohol feeding preference using the well-established Two-choice <u>Ca</u>pillary <u>Fe</u>eder (CAFÉ; SFig. 5A) assay [59], and observed that both AX and RC males exhibited significantly altered alcohol food preference compared to CV males (SFig. 5B-D). However, approximately 15% of the flies in each condition died by the end of day 5 (data not shown), and some have suggested that flies are experiencing starvation conditions in the assay [60]–[62]. This is particularly problematic for assessing alcohol preference because it is difficult to separate a preference for alcohol due to its pharmacological effects from a preference driven by its increased caloric content. To address this potential caveat to the CAFÉ results, we used BARCODE, a new starvation-independent alcohol preference paradigm that uses a sectioned stage providing unlimited food access, and promoting a more natural feeding behavior of roaming and sampling [63]. The sectioned stage contains alternating squares of solid fly food with and without ethanol in a large chamber (Fig. 4A). BARCODE permits measurement of both positional preference and food consumption preference. Positional preference is determined by counting the average number of flies on ethanol food squares versus non-ethanol food squares normalized to the total number of flies on the stage. CV and RC males had an aversion to the ethanol squares (average PI = −0.10), which decreased toward neutral by day 2 (Fig. 4B). In contrast, AX males exhibited a positional preference for ethanol that did not change significantly during the assay (average PI = +0.10; Fig. 4B,C) compared to both CV and RC males.

**Figure 4.**
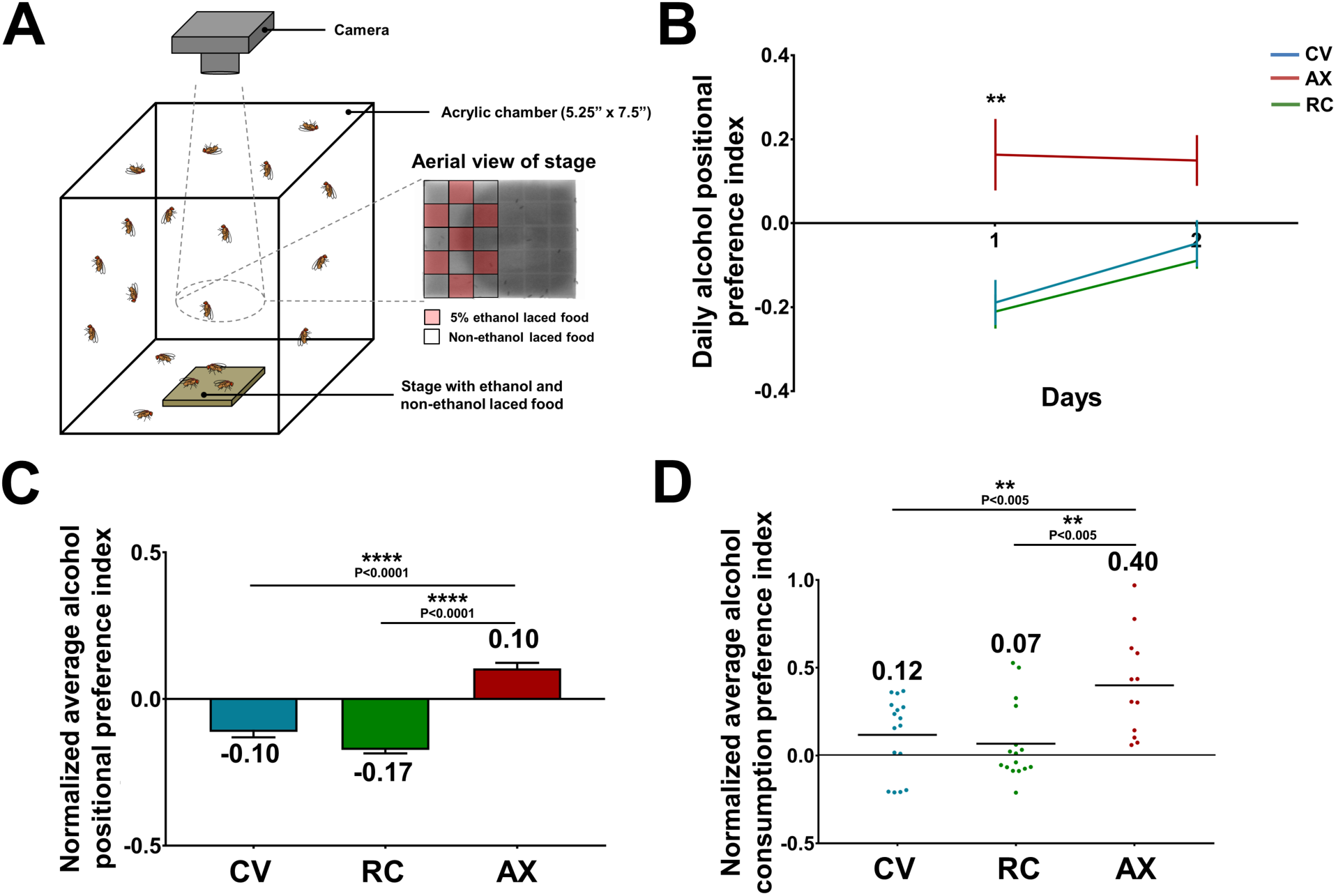
AX males exhibit a greater preference for alcohol compared to their CV or RC siblings. **(A)** A schematic of the BARCODE assay to test alcohol preference. **(B)** Daily alcohol <u>positional</u> preference for CV, AX, and RC males. On day one, AX male flies were significantly different from CV and RC flies (**, P=0.0092 and P=0.0032, respectively). No significant difference was observed on day two. n = 50 flies/condition. Bars indicate standard error of the mean and statistical significance was assigned by a two-way ANOVA (Tukey’s multiple comparisons post hoc test). **(C)** Normalized average alcohol <u>positional</u> preference index for CV, AX, and RC males over the two days of the assay. n = 50 flies/condition. Bars indicate standard error of the mean and statistical significance was assigned by a Kruskal-Wallis test (Dunn’s multiple comparisons post hoc test). **(D)** Normalized average alcohol <u>consumption</u> preference index for CV, AX, and RC males over the two days of the assay reveals that AX males have a significantly greater preference for alcohol consumption than their CV or RC siblings. Each dot represents the average of five flies. Solid black bars indicate the mean and statistical significance was assigned by a one-way ANOVA (Holm-Sidak’s multiple comparisons post hoc test).

To quantify alcohol food consumption preference, the ethanol squares and the non-ethanol squares were spiked with two different oligonucleotides. Following the two days of the assay, qPCR was used to quantify these sequences in lysates of surface-washed flies, and these values were used to calculate the alcohol consumption PI. Although CV and RC males spent less time on the ethanol squares (Fig. 4C), they consumed more ethanol food than non-ethanol food (PI = 0.12 and 0.07 respectively; Fig. 4D). Consistent with the increase in positional preference, AX males consumed significantly more ethanol food than CV or RC males (PI = 0.40; Fig. 4D). Together, the BARCODE assays indicate that AX males have a significantly stronger preference for food containing ethanol than their CV or RC siblings.

## Discussion

The microbiota’s ability to profoundly influence host physiology and behavior has important implications for understanding normal biology and disease states, and we are only beginning to discover the scope of that influence and the underlying molecular mechanisms. Using a gel-based proteomic screen, we identified both generic and sex-specific microbiota-dependent proteome changes in the *Drosophila* head. In humans, mice, and flies, the composition of the microbiota appears to be different in males and females [64]–[66], and in mice this difference is mediated by sex hormones, and sex-specific differences in bile acids and immunity [67]–[69]. The interactions between host sex and gut microbiota appear to contribute to sex-specific differences in Type 1 diabetes [67], and liver carcinogenesis [68]. While these connections are being made, there is much to learn about the mechanisms that control sex-specific differences in the microbiota and sex-specific responses to the microbiota. To date only one transcriptomic study has examined microbe-dependent gene expression changes in both male and female adult *Drosophila* tissues [32]; all other studies have focused exclusively on females [40], [70]. Thus, sex-specific microbe-dependent changes to host gene expression and protein levels have not been thoroughly investigated. We anticipate that comprehensive identification and analysis of the sex-specific difference proteins we identified in this study could yield important insight into the underlying molecular mechanisms connecting sex, the microbiota, and host biology.

One male-specific difference-protein we identified is ADH, elevated 1.8-fold in AX male heads (Fig. 2B). Because *Adh* mRNA was not responsive to microbiota elimination (Fig. 2B), we propose that ADH elevation results from increased translation or protein stability. Although ADH stability is affected by PTMs [71]–[73], we did not detect PTM changes in ADH by 2D-DIGE. Because the *Drosophila* abdominal fat body is a major site of alcohol metabolism [74], and ADH was not detectable in the brain (Fig. 2D), we propose that ADH elevation is most likely occurring in the fat body surrounding the brain. The fat body, together with the abdominal oenocytes, is functionally analogous to the liver [75], and reciprocally communicates with the brain to regulate physiology, including neural and immune activity [76]–[78]. AIH increase in AX males requires ADH activity (Fig. 3C), and ADH-dependent ethanol metabolism is required for AIH ([36] and SFig. 3A), but the precise mechanism by which ethanol metabolism mediates the elevated locomotor activity characteristic of AIH is not known. Interestingly, the lack of detectable systemic differences in ADH activity and ethanol metabolism between CV and AX flies (SFig. 3A,B) suggests that ADH elevation and activity increase in AX males is head specific. While little is known about the functional differences between head fat body and abdominal fat body, several head fat body specific GAL4 drivers exist [79], arguing for some functional distinction. Elevated ADH activity in the head fat body could protect the brain from direct effects of ethanol, such as interference with neurotransmitters [80]–[84]. Alternatively, elevated ADH could indirectly affect the brain by changing fat body metabolism and consequently altering downstream neuropeptide or immunity signaling to the brain [85]–[87].

In addition, the increase in ADH that we observe in AX males could be a mechanism to compensate for the absence of microbial-derived byproducts, like short-chain fatty acids (scFAs), that normally promote host metabolic functions [88], and affect other aspects of host development, physiology, and behavior [89]–[94]. AX flies can have abnormally high triglycerides indicative of a change in lipid metabolism [95], [96]. In flies, ADH-dependent ethanol metabolism promotes lipid synthesis directly by converting ethanol to the scFA acetate [97]. Because *Drosophila* in the wild feed on fermenting fruit (which contains up to 5% EtOH), ethanol may contribute to normal lipid metabolism [98]. Acetic Acid is also produced by the *Drosophila* microbiota and impacts development and reproductive behaviors [99], [100]. Because sex, genetic background, and nutrition affect microbiota-dependent metabolic changes [95], [96], [101], ADH levels, and physiological and behavioral responses to alcohol (this work), a comprehensive understanding of the connections between the microbiota, host metabolism, and ADH necessitates that all of these variables be taken into account. An alternative hypothesis is that the microbiota may control host ADH and physiological and behavioral responses to alcohol to improve fitness through foraging. Recent work has demonstrated that the microbiota can promote optimal foraging and enhance fitness through multiple mechanisms [13], [16], [18]. Interestingly, *Drosophila’s* responsiveness to microbe-derived alcohol and by-products of its microbe-dependent catabolism influences egg-laying behavior and promotes fitness [10].

While a microbe-dependent mechanism affecting *Drosophila’s* response to alcohol may be adaptive, varying alcohol responsiveness in humans can have harmful effects. Multiple factors contribute to the risk of alcohol use disorder (AUD) in people including disinhibition/impulsivity, patterns of alcohol metabolism, a low level of response to alcohol, and increased alcohol preference [102]. Among these, a low level of responsiveness to alcohol is the most well studied, is a strong predictor of future alcoholism, and its heritability is as high as 60% [103]. We demonstrated that AX males have an increased preference for alcohol consumption (Fig. 4), as well as increased responsiveness to alcohol as reflected in elevated AIH (Fig. 3). The sex-specific effect on responsiveness in *Drosophila* is an interesting parallel to what has been found in human studies; while decreased alcohol responsiveness is strongly associated with an increased risk of alcohol abuse in men, the same does not appear to be true in women [103]. In addition, emerging evidence suggests that alcohol consumption can cause dysbiosis of the gut microbiota observed in a subset of alcoholic patients [104], [105]. This dysbiosis appears to contribute to the neuro-inflammatory withdrawal response [106], and to the emotional effects of alcohol abuse [107]. Taken together, this accumulating evidence suggests that the microbiota may be an important contributing factor to how animals respond to alcohol. Dissecting this connection may impact our understanding of the risk factors for AUD as well.

## Author contribution

Conceptualization, B.M.M.; Methodology, M.A.B., B.M.M., J.S.M., N.S.A., and N.L.H.; Investigation, M.A.B., S.A.K., A.P., S.O., R.A.E., A.M.P., T.W.L., J.H.H., and H.M.K.; Writing – Original Draft, M.A.B.; Writing – Review & Editing, M.A.B., B.M.M., N.L.H., J.S.M., and N.S.A.; Funding Acquisition, B.M.M., J.S.M., N.L.H.; Resources, B.M.M., J.S.M., N.S.A., and N.L.H; Supervision, B.M.M. and J.S.M.

## Acknowledgements

We thank the McCartney, Minden, and Atkinson lab members for useful comments and edits during the preparation of the manuscript. We would also like to thank the Woolford, Ettensohn, Lopez, Hinman, and Linstedt labs for reagents and equipment use, and Odelia Cheng and Rebecca Lee for help characterizing the CV and RC male flies. This work was supported by a Charles E. Kaufman Foundation New Initiative Grant awarded to B.M.M., N.L.M., and J.S.M., and the UPCI Cancer Biomarkers Facility that is supported in part by award P30CA047904. A.P. was supported by NIH T32 AA07471.

## Materials and methods

### *Drosophila* stocks

Top Banana is a recent wild isolate (Seattle, WA) generously donated by M. Dickinson (CalTech). We used the following stocks from the Bloomington *Drosophila* Stock Center: Oregon R (Stock #5), Canton S (Stock #64349), and *Adh^N1^* (Stock #3976). Canton S was obtained from the Bloomington *Drosophila* Stock Center before the w-and y-transgene contamination. *Wolbachia* PCR analysis was performed on the three wild-type stocks showing that Top Banana and Canton S were infected with *Wolbachia*, and Oregon R was *Wolbachia* free (not shown).

### Creation of CV, AX, and RC cultures

CV and AX cultures were derived from embryos obtained from the same parents. A 4 hour collection of embryos was transferred in standard embryo wash (120 mM NaCl and 0.04% Triton X-100) to a plastic petri dish. For CV cultures, approximately 150 embryos were transferred to a fresh culture bottle containing autoclaved molasses fly food (8.5% molasses, 7% cornmeal, 1.1% brewer’s yeast, 0.86% agar, supplemented with 0.27% propionic acid, 0.23% methyl-parahydroxybenzoate, and 0.23% ethanol). Autoclaved inactive yeast was added to the bottle cultures (0.2-0.3g per bottle). Embryos for AX cultures were prepared as previously described [101], [102] with a few adaptations. Embryos were transferred to a separate microcentrifuge tube and treated with filtered sterilized 50% bleach for two minutes, then rinsed in sterilized 70% ethanol twice, and once with sterile water to dechorionate the embryos and eliminate microbes that were associated with the chorion. Approximately 250 AX embryos were transferred to an autoclaved culture bottle containing autoclaved yeast granules as above. The difference in embryo seeding density for the two culture conditions was necessary to ensure consistent larval density and comparable nutritional environments. All adult flies were collected 0-1 day post-eclosion into autoclaved food vials containing ~0.05g autoclaved inactive yeast granules. The flies remained in these collection vials for 5 days before being used for experiments. For RC flies, 0-1 day old AX adults were placed in preconditioned food vials that had housed ten CV males (~one week old) for 4 days. RC males were aged in preconditioned vials for 5 days. All fly cultures were reared and adult progeny maintained at 22-23°C/70% relative humidity/12-12hr light-dark cycle.

#### Verification of AX cultures

For culture-dependent verification, ten 5-6 day post-eclosion CV and AX flies were homogenized manually with a sterile pestle in 100 μL 1X PBS and three serial ten-fold dilutions were prepared. Undiluted homogenates and homogenate dilutions were then plated on MRS agar (wt/vol: 1% peptone, 1% beef extract, 0.4% yeast extract, 2% glucose, 0.5% sodium acetate, 0.1% polysorbate 80, 0.2% dipotassium hydrogen phosphate, 0.2% triammonium citrate, 0.02% magnesium sulfate, 0.005% manganese sulfate, 1% agar), Ace agar (wt/vol or vol/vol: 0.8% yeast extract, 1.5% peptone, 1% dextrose, 1.5% agar, 0.3% acetic acid, 0.5% ethanol), and Nutrient agar (wt/vol: 0.5% peptone, 0.3% yeast extract, 0.5% sodium chloride, 1.5% agar). Plates were incubated at 30°C for 48-72 hours. Plated CV fly homogenate consistently yielded robust microbial growth, while plated AX fly homogenate consistently yielded no growth (Figure S1B).

For culture-independent characterization, two 0-1 day old males were homogenized manually in filter sterilized squishing buffer (10 mM Tris pH 8.0, 1 mM EDTA pH 8.0, and 25 mM NaCl) with a fitted pestle until no obvious particulates could be seen. The homogenates were incubated with Proteinase K for 1 hour and boiled for 5 minutes. 150 ng of total DNA was used as template for PCR amplification. See Table 2 for primer information. Amplified DNAs were Sanger sequenced by Genewiz (South Plainfield, NJ).

#### Quantifying microbial load in CV and RC flies

Ten 5-6 day post-eclosion CV or RC flies were surface sterilized by sequential washes in 10% sodium hypochlorite and 70% ethanol. Flies were then washed three times with 1X PBS and mechanically homogenized in 125μL 1X PBS with ~125μL 1.0mm zirconia beads in a Mini-Beadbeater-16 (BioSpec Products) for 30 seconds. Five ten-fold serial dilutions were then prepared from fly homogenates and all dilutions were plated on both MRS and Ace agar plates. MRS plates were incubated at 37°C and Ace plates were incubated at 30°C for ~48 hours prior to counting colonies. *Lactobacillus brevis* and *Acetobacter* colonies were distinguished by characteristic colony morphologies. For each biological replicate, colonies were counted from all dilution plates with distinguishable colonies. CFU/fly values were calculated as follows: CFU/fly = ((C x D)/V) x (H / F), where C = colony counts, D = dilution factor, V = volume of diluted homogenate plated, H = volume in which flies were homogenized, and F = number of flies homogenized, as in Koyle et al. [101].

### Two-dimensional fluorescence difference gel electrophoresis (2D-DIGE), imaging analysis and protein quantification

Forty 5-6 day old flies were dry dissected on a sterile CO_2_ pad and the heads were pooled in lysis buffer (7 M urea, 2 M thiourea, 4% CHAPS, 10 mM DTT and 10 mM Na-Hepes pH 8.0) spiked with 1% protease inhibitor (Sigma-Aldrich, St. Louis MO, USA). The heads were homogenized manually with a fitted pestle until no obvious particulates could be seen. Head lysates were adjusted to 2 mg/ml protein concentration with lysis buffer using the Bradford standardizing method. Protein lysate solutions containing a total of 100 μg of protein were labeled with 2 μl of either 1mM propyl-Cy3-NHS or 0.83 mM methyl-Cy5-NHS (CyDye DIGE Fluors; GE Healthcare) as described previously [103], resulting in fewer than one dye molecule per protein to prevent changing protein migration in 2DE gels. Reciprocal labeling experiments were performed to control for dye-dependent changes and to generate technical replicates of each sample. Two-dimensional electrophoresis (2DE) was performed as previously described [104]. After second dimension electrophoresis, the gels were fixed in a solution of 40% methanol and 10% acetic acid overnight then imaged in a lab built imager [32]. Protein differences were determined by quantifying grey scale images of each channel using Source Extractor as previously described [32], [33]. To determine the fold-difference between CV and AX expression of a protein, the intensity values of each channel were normalized to five “guide star” proteins, protein spots that reliably do not change within the proteome (determined from multiple biological replicates), and analyzed as previously described [28], [32], [105].

### Immuno-blotting of ADH protein

For standard western blots, CV brain and head capsule lysates were prepared in 2X Laemmli sample buffer treated with 0.1% Protease Inhibitor Cocktail (Sigma-Aldrich, St. Louis MO, USA) and separated by SDS-PAGE. For 2D-Westerns, CV head lysate was labeled with Cy5 dye and run on a 2DE gel, as described above. Gels were equilibrated in a Tris buffer at pH 7.5 with 20% glycerol for 30 minutes. Proteins were transferred to Protran nitrocellulose membranes (Whatman, Little Chalfont Buckinghamshire, UK) overnight in carbonate transfer buffer at pH 9.9 (100mM NaHCO_3_ and 80uM Na_2_HCO_3_) at constant 25 V. Membranes were immuno-blotted using a goat anti-*Drosophila* ADH antibody at 1:500 (Santa Cruz Biotechnologies, Dallas TX, USA). Donkey anti-goat HRP secondary antibody (Jackson Immunoresearch, West Grove PA, USA) was used at 1:2,000. Chemiluminescence (Pierce ECL Western Blotting Substrate, Thermo Scientific) was detected using a ChemiDoc MP Imaging System (Bio-rad).

### qRT-PCR analyses of *Adh* gene expression levels

Five to six day old CV and AX fly heads were dry dissected under CO_2_ sedation. Ten heads per sample were immediately bead beaten with approximately 50μl of 0.5 mm Zirconia/Silica beads in 500μl of Trizol (Invitrogen, Carlsbad CA, USA). The homogenates were immediately frozen at −20°C until RNA extraction (stored no longer than 2 weeks).

RNA was extracted from *Drosophila* heads with the Direct-zol RNA mini kit (Zymo, Irvine CA, USA). 500 ng of high quality RNA (A_260/280_ ~1.8-2.0) was used as template for the synthesis of first strand of cDNA using the SuperScript VILO synthesis kit (Invitrogen, Carlsbad CA, USA). After first strand cDNA synthesis, 100 ng of the cDNA product was directly used for qRT-PCR using the PowerUp SYBR Green Master Mix in a 7300 Real Time PCR System (Applied Biosystems, Foster City CA, USA) using Sequence Detection Software (v1.4.0.25, Applied Biosystems, Foster City CA, USA).

Housekeeping gene *rpl32* (*ribosomal protein L32*) was used for normalization [106], and data were analyzed using the Pfaffl-ΔΔCT method [107] in Fig. 2C. We determined the primer efficiencies for all primer sets used to calculate the fold-change between CV and AX flies. Fold changes presented are the mean results from six biological replicates for males and four for females.

### Assessing alcohol induced hyperactivity

Locomotor activity was analyzed with a *Drosophila* Activity Monitor 2 (DAM2; Trikinetics, Waltham, MA, USA) that can accommodate a total of 32 flies in individual tubes (Fig. 3A). For inhibitor conditions, CV and AX male flies were aged for five days in vials with sterile fly food and sterile yeast paste containing 5μM 4-methylpyrazole (Sigma-Aldrich, product code: 286672). For all experiments using the DAM2, single 5-6 day old adults were placed under mild CO_2_ sedation and transferred into monitoring plastic tubes (5 mm diameter) capped at one end with a rubber cap. Tubes were randomly loaded onto a Trikinetics exhaust manifold. Base line activity was established for the first hour followed by a 2 hour ethanol exposure (PHARMCO-AAPER, Shelby KT, USA) using a mixture of air to ethanol vapor at a 10:1 ratio (empirically determined for max hyperactivity difference) delivered by a homemade vaporizer. Activity data (crossings of an infrared beam) were collected by the Trikinetics computer software in bins every 30 seconds and later converted to 5 minute bins. A total of 8 flies from each condition were run in parallel in any given experiment. Data were pooled from different trials for each condition to obtain average activity levels. Locomotor activity curves were generated and statistical analysis was performed with GraphPad Prism 7. A two-way ANOVA was used to compare the hyperactivity curves.

### Assessing alcohol induced sedation

The alcohol sedation assay was performed as described in Maples and Rothenfluh (2011) with minor adaptations. 0-1 day adult males were collected in batches of 8 under CO_2_ and aged to 5-6 days. Fresh fly food vials were converted into ethanol chambers by creating a flat cotton bed at the bottom of the vial and sealing the chamber with a cotton ball soaked in 100% ethanol [50] (PHARMCO-AAPER, Shelby KT, USA). The chamber size was approximately 1.25” in height.

A typical alcohol sedation experiment contained 4-5 vials each of CV males and AX males. The flies were transferred to ethanol chambers, conditions randomized, and numbered. The vials were taped together in batches of 4-5 vials. An iPad was used to record videos of each trial. Before recording, each dry ethanol chamber cotton ball was replaced with a cotton ball soaked in roughly 1.2 mL of red dyed ethanol. Time zero started after all dry cotton balls were replaced. During the experiment, the vials were tapped every minute and immobility was assessed after a 15 second recovery period until full sedation was reached. After the 15 second recovery period, the number of flies immobilized was recorded and the mobile fraction was calculated. The ethanol cotton ball was readjusted after every tap series to ensure the chamber was approximately 1.25” high throughout the experiment. The videos were analyzed by two observers who were blinded to the microbial conditions of the flies. Flies were deemed immobile if: (1) the fly traveled less than the radius of the vial (to account for postural struggle or spontaneous jumps after immobilized), (2) the fly lost postural control and flipped orientation, or (3) the fly was completely motionless or stationary with tremors while maintaining postural control. Statistical significance was determined by performing a two-way ANOVA comparing the sedation curves and a one-way ANOVA for the ST50 values using GraphPad Prism 7.

### Assessing alcohol food preference using the Two-choice Capillary Feeder (CAFE) assay

CV, AX, and RC male sibling flies were tested in the CAFE assay as previously described [54]. Each vial housed 8 flies and contained 4 capillaries. The capillaries contained a liquid food comprised of 5% yeast extract and 5% sucrose with either no ethanol (2) or 10% ethanol (2). Measurements of food levels in the capillaries were taken daily by four observers. Death counts for each vial were also noted per day. The assay was carried out for 5 days and measurements were taken at the same time each day. Capillaries were replaced with fresh food solution each day.

### BARCODE alcohol preference assay

Fifty 5-6 day old CV, AX, RC flies were tested in parallel in three separate BARCODE chambers. Standard molasses based fly food with agar was used for preference assays. Fly food was liquified and additives were mixed in once the food was cooled to ~35 °C. The food grid was filled in an alternating pattern with food containing 5% ethanol or non-ethanol food to which a matching volume of water was added. The food type specific oligomers were added to the corresponding food type at 3.5 ng/μl. After 2 days, the flies were collected from the chamber using CO_2_ and frozen immediately for future DNA extraction (see below).

#### Positional alcohol preference analysis

Preference was tested behaviorally in the BARCODE assay by capturing images of the position of flies on the food pad in 5 minute intervals for 48 hrs. We used BTV Pro for Mac (Ben Software) for automatic capture and analyzed using a custom Perl script and ImageMagick (ImageMagick 7.0.5-0). The Preference Index (PI) was measured by counting the number of flies on ethanol and non-ethanol food squares per day and calculated as PI = (N Flies on Ethanol - N Flies on Non-Ethanol) / Total N Flies on Stage.

#### Consumptive alcohol preference analysis

Consumptive preference was measured following the conclusion of the behavioral assay. Flies were washed using the washing protocol in [108] and homogenized in a squishing buffer (10 mM Tris-HCl @ pH 8.2, 1 mM EDTA, 25 mMM NaCl). For each qPCR biological replicate, we homogenized five flies per sample (n=5). Homogenates were then incubated with Proteinase K (NEB, Ipswich, MA, Product No. P8107S) and spun down for 2 minutes at 10,000 G. Ten microliters of the supernatant was used for qPCR with the ThermoFisher Power SYBR™ Green PCR Master Mix reagents (Waltham, MA, Catalog No. 4367659) and analyzed with a ThermoFisher Viia 7 Real-Time PCR System (Waltham, MA) with a T_m_ = 60 °C and 40 cycles per run.

### Measuring alcohol levels

20 males per biological replicate were pretreated with 10% ethanol vapor for 30 minutes with a homemade vaporizer then frozen immediately. Flies were homogenized with glass dounce homogenizers using 20 μL ddH_2_O per fly. The homogenate was pipetted into 1.5 mL tubes and centrifuged at 10,0000 G for 3 minutes. A 10 μL aliquot of the supernatant was assessed using the Megazyme Ethanol Assay Kit (Bray, Co. Wicklow, Ireland, Product code: K-ETOH). Absorbance measurements were taken using a Nanodrop ND-1000 (Nano-drop Technologies, Inc., Wilmington, DE). The A_1_ measurement was taken after 5 minutes following the addition of the ALDH enzyme and the A_2_ measurement 15 minutes following the addition of the ADH. ALDH absorbance was background subtracted from ADH absorbance to calculate NADH produced.

### Measuring alcohol dehydrogenase enzymatic activity

The Alcohol Dehydrogenase activity assay kit from Sigma-Aldrich (product code: MAK053) was used. Twenty males per biological replicate were pretreated with 10% ethanol vapor for 30 minutes with a homemade vaporizer then immediately homogenized with a fitted plastic pestle in 200 μL of ice cold ADH assay buffer, in a 1.5 mL microcentrifudge tube, until no particulates were visible. The homogenates were centrifuged at 10,0000 G for 10 minutes. Ten μL supernatant samples were assessed for ADH activity. Absorbance measurements were taken using a Tecan Safire 2 Plate Reader (Tecan Group Ltd., Männedorf, Zürich, Switzerland) and from these an NADH standard curve was generated. The initial absorbance measurement was taken two minutes following the addition of the enzyme mix (ADH assay buffer, developer, isopropanol substrate). The reaction was run at 27°C and absorbance measurements were recorded every minute for a total of 25 minutes. A ∆Absorbance (∆A) value was calculated for each sample by subtracting the initial absorbance from the final absorbance value. The ∆A was used to calculate the ADH activity using the following equation:

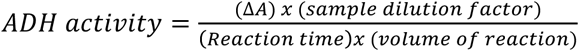

### Statistical analysis

All analyses were performed with GraphPad Prism 7 (GraphPad Software, Inc., La Jolla, CA). For comparisons with only two conditions, a Student’s t test (or Mann-Whitney for non-normal distributed data) was performed. For experiments containing three or more conditions, an ANOVA analysis was used. Various posthoc analyses were performed depending on the type of comparison and sample distribution. For comparisons where experimental conditions were compared to only the control, a Dunnett’s comparison was used. When all conditions were compared to each other and all samples had normal distributions, a Tukey’s comparison was used. When all conditions were compared to each other, and when at least one condition had a non-normal distribution, a Sidak’s comparison was used.

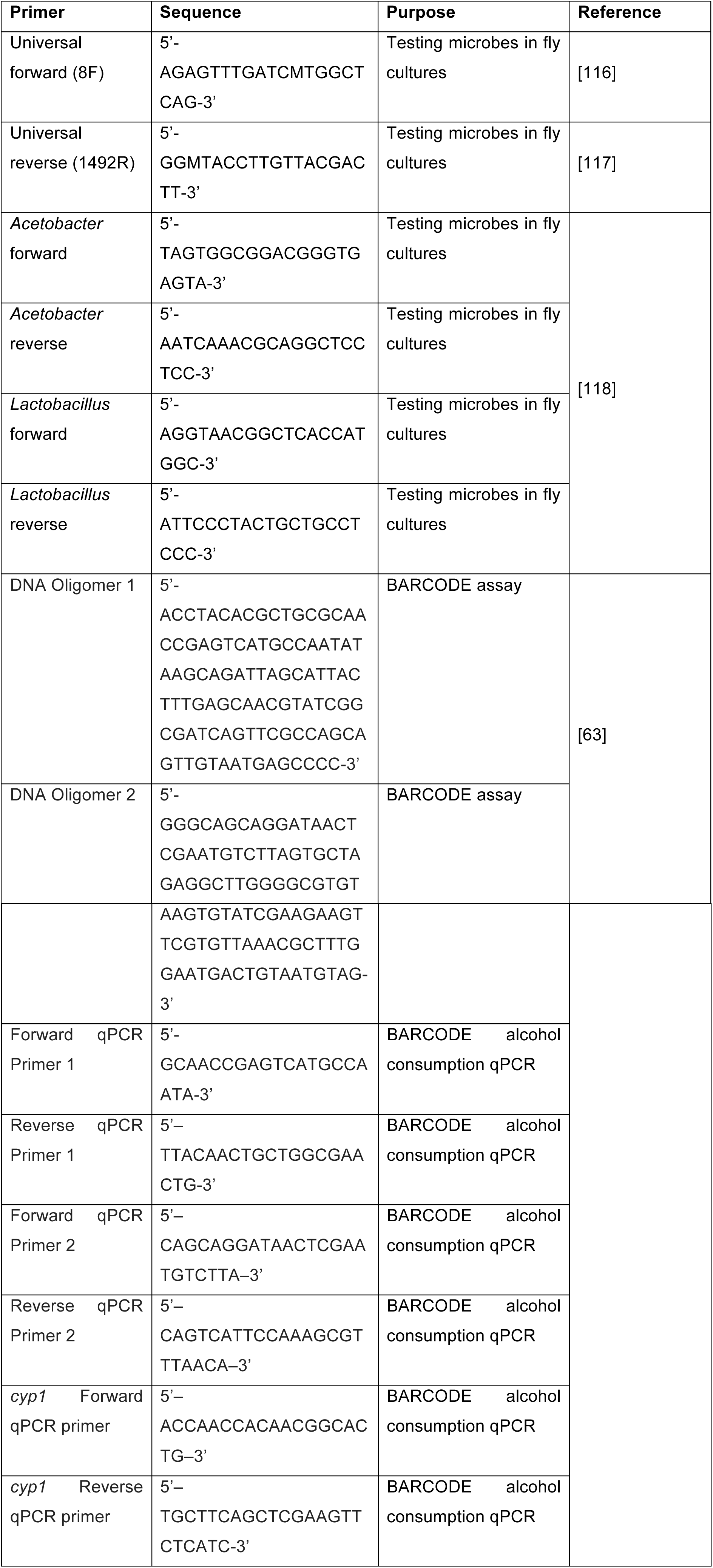
Primers used in this study.

